# Temporal Dissociation of Syntactic Disambiguation and Memory Retrieval during Sentence Processing: Naturalistic MEG Evidence from Interpretable Models

**DOI:** 10.64898/2026.04.20.719609

**Authors:** Donald Dunagan, Dylan Scott Low, Shisen Yue, Lars Meyer, John T. Hale

## Abstract

Human sentence comprehension proceeds word-by-word, with prior research proposing two central sources of cognitive demand during incremental processing: forward-looking disambiguation of the incoming information stream, and backward-looking retrieval of information associated with previous words from working memory. Recent work has shown that Transformer-based language models successfully generate predictions about sentence processing load in human psycho- and neurolinguistic data by operationalizing disambiguation cost as next-token surprisal, and memory retrieval cost as normalized attention entropy (NAE). Such models, however, remain difficult to interpret as it is not well understood what factors play causally into the decision to assign a cost value to a given word in such artificial neural networks. Here, we present interpretable and cognitively grounded models of disambiguation and memory retrieval and evaluate their neural alignment and spatio-temporal correlates using human magnetoencephalography responses to naturalistic narrative speech. Multivariate temporal response function modeling demonstrates firstly that these human-bias-informed models fare equally well in accounting for observed human language processing data as their Transformer counterparts. This same modeling framework then suggests that surprisal and NAE temporally dissociate in the cortical language network — surprisal being predictive of bilateral superior temporal gyrus and supramarginal gyrus activation ∼300–500 ms, and NAE being predictive of activity in the same regions, but later ∼750–850 ms. By demonstrating that interpretable neurocomputational models can achieve meaningful brain alignment while maintaining explanatory transparency, this work offers a methodological blueprint for bridging the gap between algorithmic theory and neural implementation.

## Introduction

Sentence comprehension involves deriving an understanding from an incoming stream of linguistic information. In order to perform this incremental cognitive task, however, the human sentence processor must both be able to handle ambiguity, as well as have a working memory for the storage and later retrieval of perceived information. Thus, two central sources of cognitive demand during sentence processing are disambiguation and memory retrieval. Disambiguation involves arriving at the correct analysis when multiple alternative analyses are consistent with the input encountered thus far. With each structural and/or lexical analysis being associated with a probability, word-by-word processing difficulty can be quantified via information-theoretical complexity metrics (e.g., surprisal: Hale, 2001; R. Levy, 2008; entropy reduction: Hale, 2003, 2006, 2011; see Hale, 2016 for a review). Such “rightward-looking” processes are only one half of the story, however; the other being memory demand — which may be considered “leftward-looking.” One influential theory of memory demand is cue-based retrieval (Lewis and Vasishth, 2005; Lewis et al., 2006; Van Dyke, 2007; Van Dyke and Lewis, 2003; Van Dyke and McElree, 2006, 2011, inter alia; see Vasishth and Engelmann, 2021 for a monograph-length discussion and review). Under this account, an incoming word triggers a retrieval operation over information associated with previous words in order to resolve grammatical dependencies between the current word and the preceding context.

Research has found that word-by-word metrics — of both disambiguation and memory retrieval difficulty — derived from neural language models (LMs) are predictive of data recorded during human sentence processing (disambiguation: Frank, 2013; Frank et al., 2015; Hao et al., 2020; Heilbron et al., 2022; Oh and Schuler, 2023; Pimentel et al., 2023, inter alia; memory retrieval: Oh and Schuler, 2022; Ryu and Lewis, 2021, 2025; Yoshida et al., 2025). Yet, neural LM accounts — owing to their diminished degree of interpretability — shed little light on the mechanisms proposed to underlie human sentence comprehension. Here, we present interpretable, cognitively grounded models of disambiguation and memory retrieval and demonstrate: 1) that they fare just as well in accounting for observed human language processing data as their neural LM counterparts; and 2) that they, when combined, elucidate components of a staged processing (Sternberg, 1969; cf. McClelland, 1979) account of incremental^1^ sentence processing (Bornkessel & Schlesewsky, 2006; Forster, 1979; Frazier & Fodor, 1978; Friederici, 2002) in which syntactically-informed lexical disambiguation occurs prior to cue-based retrieval from working memory.

We use a publicly available magnetoencephalography (MEG) dataset collected by Brodbeck et al. (2022) to probe the spatio-temporal correlates of disambiguation and memory retrieval during naturalistic language comprehension. Following Gwilliams et al. (2025), we distinguish cognitive operations from cognitive functions. We treat both disambiguation and memory retrieval as operations subserving the overall function of syntactic-level sentence processing. That is, in terms of Marr’s (1982) tri-level framework for cognitive science, disambiguation and memory retrieval are algorithmic-level proposals. By probing the neurological and temporal loci of these algorithms via naturalistic MEG, we thus seek to bridge the gap between algorithmic-level proposals and implementation-level evidence.

With an eye toward the twin goals of both explainability and neurocomputational alignment, in the present study, we evaluate two interpretable models against the much less transparent GPT-2 (Generative Pre-trained Transformer; Radford et al., 2019). As an algorithmic theory of disambiguation, we consider surprisal derived from the dependency parser introduced by Dunagan (2025, Subsection Incremental Left-Corner Generative

Dependency Parsing). In regard to memory retrieval, we consider normalized attention entropy derived from the cue-based retrieval model of Yue (2026, Subsection Grammar-Bilinear Asymw2v) — a specific implementation of Van Dyke’s (2007) theory of interference effects. We use the multivariate temporal response function modeling technique (Brodbeck et al., 2023; David et al., 2007; Lalor et al., 2006, Subsection Multivariate Temporal Response Function Modeling) for the neurocomputational modeling of naturalistic MEG data.

### Incremental Left-Corner Generative Dependency Parsing

Language models calculate probabilities over word strings, where the probability of a sequence of words *p*(*w*_1:_*_n_*) = *p*(*w*_1_*, w*_2_*,…, w_n_*) can be calculated using the chain rule of probability:

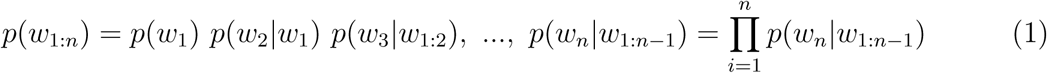

Whereas GPT-2-caliber LMs make use of the Transformer neural network architecture (Vaswani et al., 2017) and semi-large-scale pre-training^2^ to learn such probabilities, LMs can also be designed and trained with alternative inductive biases in mind. Here, we focus on one such model, the Incremental Left-Corner Generative Dependency Parser (ILCGDP) presented in Dunagan (2025, Ch. 2). In doing so, we take parsing — i.e., the assignment of grammatical structure to an inputted sentence string — to be a fundamental component of disambiguation during human sentence processing. Indeed, it has been demonstrated neurally (Brennan et al., 2016, 2020; Hale et al., 2015; Shain et al., 2020; Wolfman et al., 2024), electrophysiologically (Brennan & Hale, 2019; Hale et al., 2018), and in self-paced reading (Fernandez Monsalve et al., 2012), that surprisal derived from syntactically-informed language models can improve model quality or better fit human language processing data than surprisal derived from syntactically-uninformed language models.

The ILCGDP operates as a dependency parser, mapping an input sentence to a dependency graph under some dependency grammar — a formal framework for the notion of linguistic dependency (de Marneffe & Nivre, 2019; Gaifman, 1965; Hays, 1964, 1965; Robinson, 1967; Tesnière, 1959). As compared to familiar constituency-based phrase structure grammars, in which elements hierarchically combine to form constituents (phrases), in a dependency grammar, the words of a sentence relate to one another via binary asymmetric (directed) dependencies, which are commonly, but not necessarily, typed, i.e., labeled. Each dependency has a head, or governing word, and a dependent (child), or governed word. The constraint is often imposed that each word may have only a single head. The main verb of a sentence serves as the head of the sentence and can be thought of as being governed by an artificial utility element ROOT (Ballesteros & Nivre, 2013). From the position of the main verb, all other words in the sentence can be reached by traversing dependencies. Sentence-final punctuation, when it is included, is governed by the main verb. It is this collection of dependencies for a sentence — here, assigned by the ILCGDP — which make up a dependency graph and the sentence’s assigned grammatical structure.

Fig. 1 gives a labeled dependency graph for example (1).

**Figure 1.**
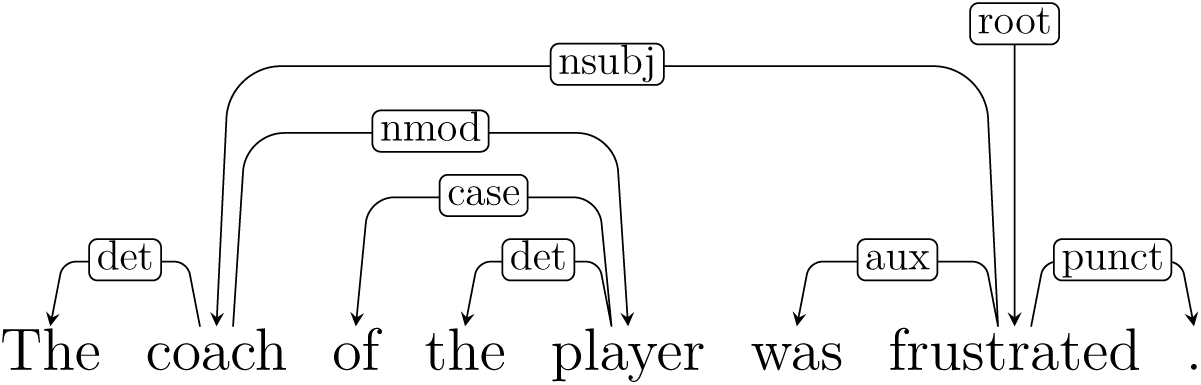
Dependency graph for example (1).

(1) The coach of the player was frustrated.

Dependency graphs draw dependencies as directed edges (arcs; from head to dependent), often labeled with the tag of the relation. For example, in Fig. 1, the word *frustrated* is labeled by root because it is the main verb; *frustrated* in turn governs *coach* via an nsubj (nominal subject) relation, as well as *was* via an aux (auxiliary) relation.

The ILCGDP (Fig. 2) is of interest here because it is a grounded and transparent model of disambiguation, assigning labeled dependencies to natural language in a manner which incorporates cognitively plausible biases: 1) it operates incrementally, with only marginal lookahead; 2) it is generative, modeling both explicit syntactic structure as well as word probability; and 3) it is left-corner, making predictions about grammatical relations for as of yet unseen words, and operating with working memory demands that mimic those of the human sentence processor.

**Figure 2.**
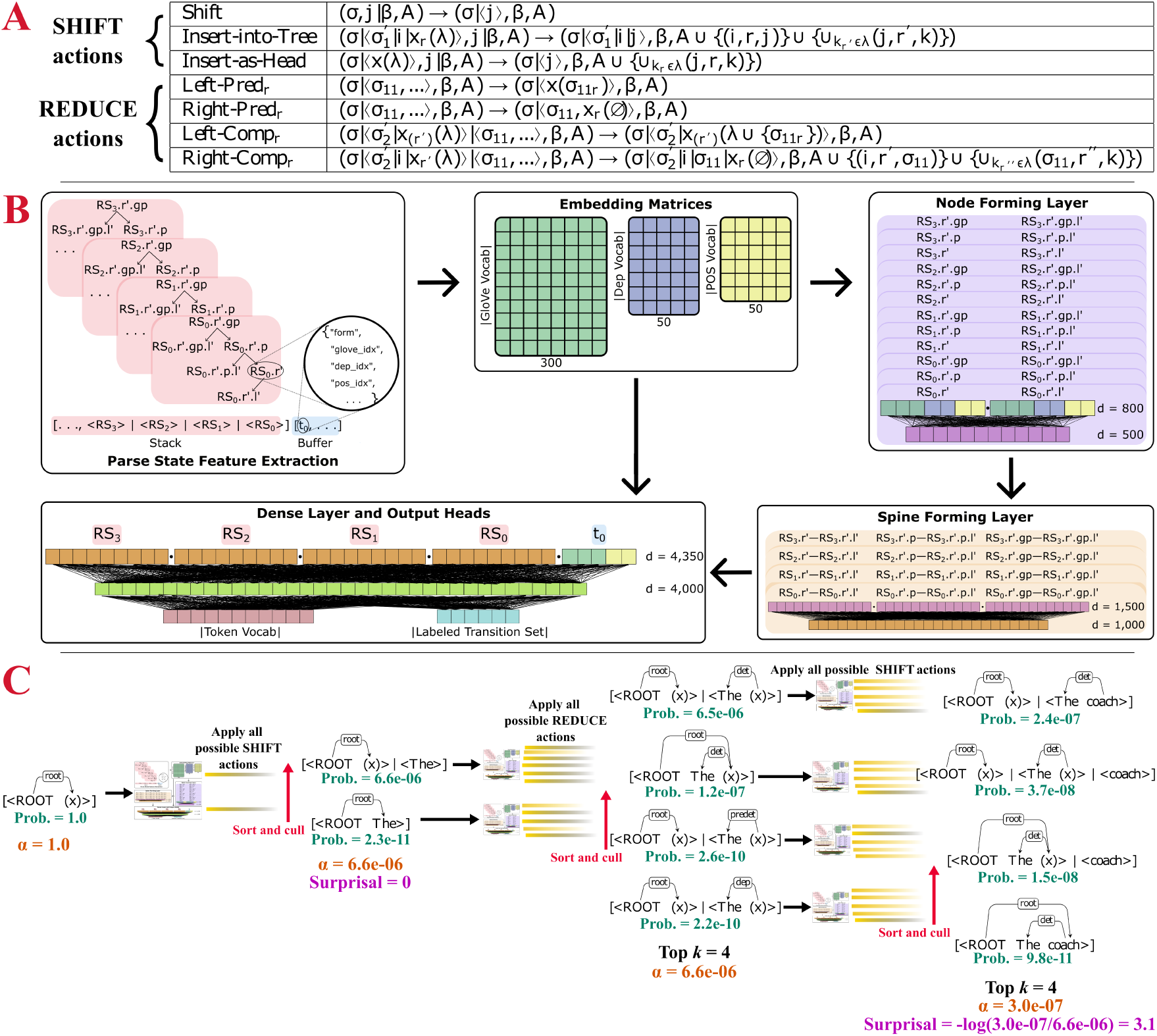
Aspects of the Incremental Left-Corner Generative Dependency Parser. (A) The transition set describes structure building and information processing rules, and how those rules update the present state of affairs. (B) Feature template demonstrating how numerical representations are extracted from the linguistic-symbolic parse state and the architecture of the deep neural network model used to assign scores to transitions and upcoming words. (C) Demonstration of beam search parsing and surprisal calculation. At each parsing step, the k probability-ranked partial parses on the beam are each passed through the trained neural network to generate all possible states reachable by one transition. This collection of configurations is sorted and then culled to just the top k once again, based upon probability. α indicates prefix probability, or sum of the probabilities of the partial parses on the beam. Surprisal of the second word (and those subsequent) is calculated as the negative log ratio of the current to former prefix probability for beams resulting from SHIFT actions.

### Incremental Transition-Based Dependency Parsing

Like an autoregressive language model — and a human sentence processor, for that matter — the ILCGDP operates in a left-to-right manner, with a single pass through the input string.^3^ Indeed, it is an instantiation of transition-based dependency parsing (Kudo and Matsumoto, 2002; Nivre, 2003; Yamada and Matsumoto, 2003; see Kübler et al., 2009; Nivre, 2006 for reviews). Transition-based dependency parsing implements an abstract machine in which a configuration represents the current state of affairs, while collections of rules describe transitions between configurations which build structure and process incoming information. Two special states are defined: an initial state and a terminal state. In the initial state, no structure has yet been built and all of the input information remains ready to be processed; in the terminal state, a complete dependency parse has been constructed and the entirety of the input string has been read in. In this paradigm, the parsing task is the selection of a good sequence of transitions, which traverse from the initial state to the terminal state while constructing the correct dependency parse.

### Left-Corner Parsing

Building off of the work of Noji and Miyao (2014, 2016), the ILCGDP implements a left-corner transition system. Classically, transition-based dependency parsing operates bottom-up (Kuhlmann et al., 2011; Nivre, 2003, 2004; see also Johnson-Laird, 1983 and Aho and Ullman, 1972), with nodes (tokens) — owing to a lack of non-terminal nodes that would be predicted top-down — being enumerated as they are read from the input string, and with dependencies only capable of being established once the associated governing and governed words are both available for consideration (see Nivre, 2004, p. 6).

Related to the observation that bottom-up phrase structure parsing has space (memory) complexity which grows proportionally with right-branching depth (Abney & Johnson, 1991; Resnik, 1992), Noji and Miyao (2014, 2016) show that the traditional bottom-up dependency parsing transition sets cannot guarantee constant space complexity for right-branching constructions. This memory demand profile is unaligned with human sentence processors, who have no difficulty processing right-branching structures (Miller & Chomsky, 1963). A more cognitively realistic transition system for dependency parsing is thus motivated.

With the aim of developing a dependency parser with constant memory demands for left- and right-branching structures, but non-constant memory demands for center-embedding structures — in line with human processing difficulty (Abney & Johnson, 1991; Miller & Chomsky, 1963; Resnik, 1992) — Noji and Miyao (2014, 2016) draw inspiration from the left-corner parsing automaton of Resnik (1992) to develop a left-corner transition set for dependency parsing which combines true top-down prediction with bottom-up composition. In order to accommodate the idea of predicted nodes as in left-corner constituency parsing, they introduce dummy nodes, which serve as subtrees to hold the place of words which are yet to be processed.

The parser utilized here — the ILCGDP — was presented in Dunagan (2025) and further develops this framework in two significant ways: 1) it modifies and expands upon the Noji and Miyao (2014, 2016) transition set so that the dependencies assigned between words can be labeled (e.g., subject, direct object, conjunction, adverbial modifier; the complete ILCGDP transition set is given in Fig. 2A); and 2) it is generative, modeling both explicit syntactic structure as well as word probability.

### Generative Modeling

A generative parsing model is one which defines a joint distribution on words **x** and parses **y**: *p*(**x**, **y**). Transition-based models — like the ILCGDP — are amenable to generative modeling, simply requiring that: 1) transitions are scored probabilistically, given a configuration (see Fig. 2B); 2) the probabilities for transitions which generate the next word are multiplied by the probability of generating the next word, given a configuration; and 3) forward lookahead is restricted such that information “from the future” cannot be used to make decisions regarding the set of circumstances in the present. Because the parser state in transition-based parsing encodes information about the collection of previously applied transitions — i.e., it is a form of history-based parsing (Black et al., 1993) — the chain rule of probabilities can be applied to decompose the probability of a sentence *x* and a parse *y* into a sequence of actions (*a*_1_, *…*, *a_T_*), where *a_T_* is the action that completes the tree:

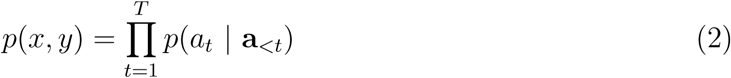

While surprisal — our present operationalization of disambiguation — is calculated from beads-on-a-string language models such as GPT-2 as negative log conditional probability (R. Levy, 2008):

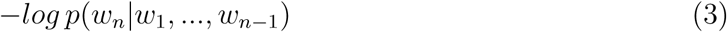

surprisal from generative models like the ILCGDP is calculated as the log ratio of the prefix probabilities before and after a word, where the prefix probability at a word is the sum over the probabilities of compatible structures at that word (Hale, 2001, see Fig. 2C):

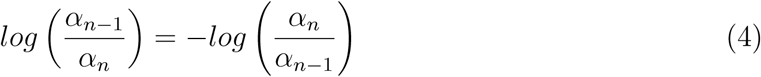

It should be noted that parsers like the ILCGDP that are guided by a generative model face an imbalance regarding the probabilities of lexical and structural actions. This imbalance is well known in natural language processing (Buys & Blunsom, 2015a; Crabbé et al., 2019; Fried et al., 2017; Stanojević & Steedman, 2020; Stern et al., 2017) and carries consequences for neurobiology (see Hale et al., 2018 for EEG or Franzluebbers et al., 2024 for fMRI). It arises because the probabilities of word generation actions are so much smaller than the probabilities of dependency forming actions. Concretely, the ILCGDP has available to it a total of 179 structural operations — a stark contrast to the output vocabulary of 39,552. Thus, when calculating word-by-word surprisal, the change in beam prefix probability which is attributable to syntactic structure building is minuscule in comparison to the change attributable to next token generation. So while the predictor combines both types of disambiguation, it is the syntactically-informed disambiguation of the next word choice that shows up most strongly in the derived complexity metric. This sort of disambiguation is roughly comparable to that performed by opaque language models (i.e., GPT-2).

### Grammar-Bilinear Asymw2v

Apart from disambiguation, we also take word-by-word language comprehension to involve retrieval from working memory for the establishment of dependencies with the current word. Cue-based retrieval theory proposes: 1) that memory access is content-addressable, with morphological, syntactic, and semantic cues from the current word probing working memory for compatible candidates; and 2) that partially matching distractors are responsible for similarity-based interference effects (Lewis et al., 2006; Van Dyke & McElree, 2006). Recent work has drawn an analogy between cue-based retrieval and diffuseness of self-attention in neural LMs (Oh & Schuler, 2022; Ryu & Lewis, 2021, 2025; Timkey & Linzen, 2023; Yoshida et al., 2025) — the more diffuse an attention pattern, the greater the interference inherent in a retrieval operation (caused by the presence of distractor items similar to the true target).

We operationalize memory retrieval burden as normalized attention entropy (NAE; Oh & Schuler, 2022). For a sequence of words *w*_1_*, w*_2_*,…, w_n_*, where *w_n_* is the word currently being processed (i.e., the one generating the retrieval cue), NAE is calculated based on attention scores assigned to all of the preceding words *w*_1_*, w*_2_*,…, w_n−_*_1_. While previous work has used attention scores from GPT-2 (Oh & Schuler, 2022; Ryu & Lewis, 2021, 2025), we use the Grammar-Bilinear Asymw2v (“asymmetric word to vector;” GBA) model presented in Yue (2026). This model was developed from the outset: 1) with transparency in mind; and 2) to consider the roles played by both syntax as well as semantics in memory retrieval (Van Dyke, 2007; Van Dyke & McElree, 2011).

### Normalized Attention Entropy

The NAE complexity metric is an extension of the attention entropy complexity metric proposed by Ryu and Lewis (2021, 2025). Interpreting the self-attention mechanism of the Transformer architecture as implementing cue-based retrieval of the sort theorized by Lewis et al. (2006), attention entropy is intended to measure similarity-based interference during such retrievals, with the presence of distractor items similar to the true target causing a greater spread in the self-attention pattern. In Ryu and Lewis (2025), the attention entropy at word *w_n_* is defined as:

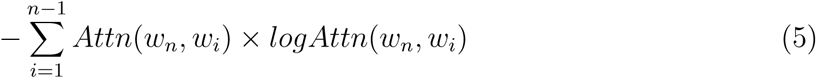

This “vanilla” attention entropy metric, however, suffers the drawback that entropy systematically tends to be higher over longer sequences. It is thus confounded by sentence length, meaning that sentences of different lengths are not directly comparable.

Furthermore, Ryu & Lewis attention entropy operates on improperly normalized probability distributions, as attention weights over previous tokens *w*_1_*,…, w_n−_*_1_ sum to less than 1. NAE corrects for both problems by: 1) re-normalizing the attention weights over the previous tokens *w*_1_*,…, w_n−_*_1_ to sum to 1; and 2) calculating the observed attention entropy proportionate to the maximum possible entropy at *w_n_* (Oh & Schuler, 2022):

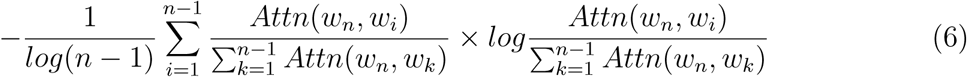

Fig. 3 provides a graphical demonstration of attention diffuseness in the context of example (1). At the word *frustrated*, a retrieval is made for the subject of the sentence in order to form the nsubj dependency relation. As an aside, in the Universal Dependencies framework (UD; de Marneffe et al., 2014, 2021; Nivre et al., 2016) used by the GBA for triggering retrieval operations, the nsubj dependency relation governing the nominal subject originates from the main verb, rather than from the auxiliary (see Fig. 1). Regardless, a portion of attention is pulled away from the true subject (*coach*) by the distractor (*player*). Under the present proposal, this attention entropy is expected to be correlated with similarity-based interference.

**Figure 3.**
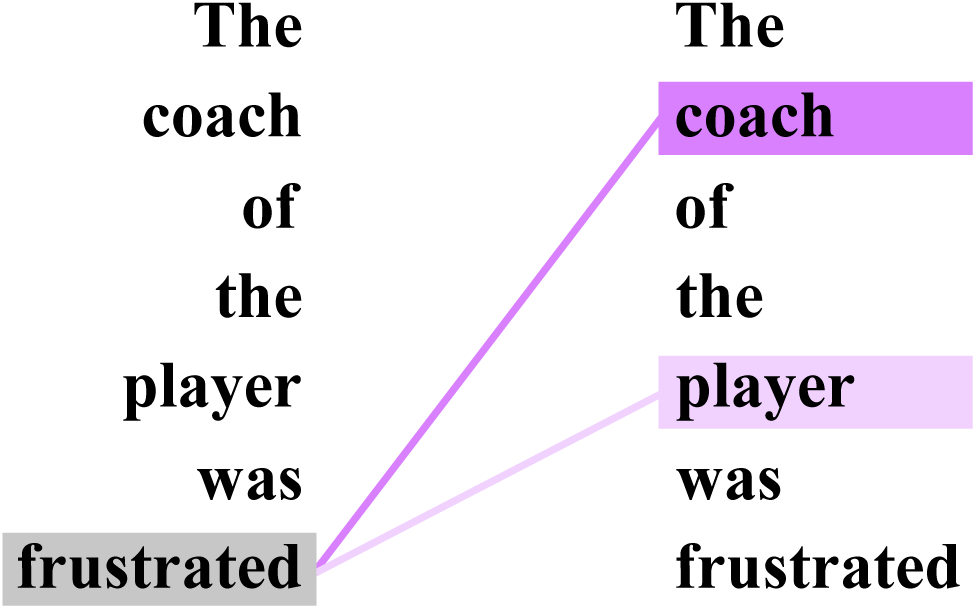
Potential distribution of attention at the word frustrated — from example (1) — when making a retrieval to form the nsubj dependency relation with the subject of the sentence (see Figs. 1 and 4).

### The Mixed Grammar-Bilinear Asymw2v Model

The GBA (Fig. 4) is a lightweight linear model that factors syntactic and semantic contributions to interference while staying close to the retrieval computation, itself. The approach treats dependency arcs (following Hofmeister and Vasishth, 2014; Van Dyke and McElree, 2006) as retrieval traces and models retrieval as an asymmetric linear transformation from a cue representation into the target’s representational space. Additional details immediately follow, but in brief: 1) the syntactic submodel (the grammar-bilinear component) uses Combinatory Categorial Grammar supertags (CCG; Steedman & Baldridge, 2011) and a bilinear parameter *W* to score cue–target compatibility and capture similarity-based syntactic competition; and 2) the semantic submodel (the asymmetric word to vector component) adapts dependency-based embeddings (O. Levy & Goldberg, 2014) to learn distinct input (cue) and output (relation-bound target) spaces, optimizing proximity for words and their dependency-labeled contexts, while repelling unrelated items. By separating these two sources of competition within a linear retrieval framework, the model provides an interpretable parametrization of interference that can be analyzed directly in terms of syntactic structure and lexical compatibility.

**Figure 4.**
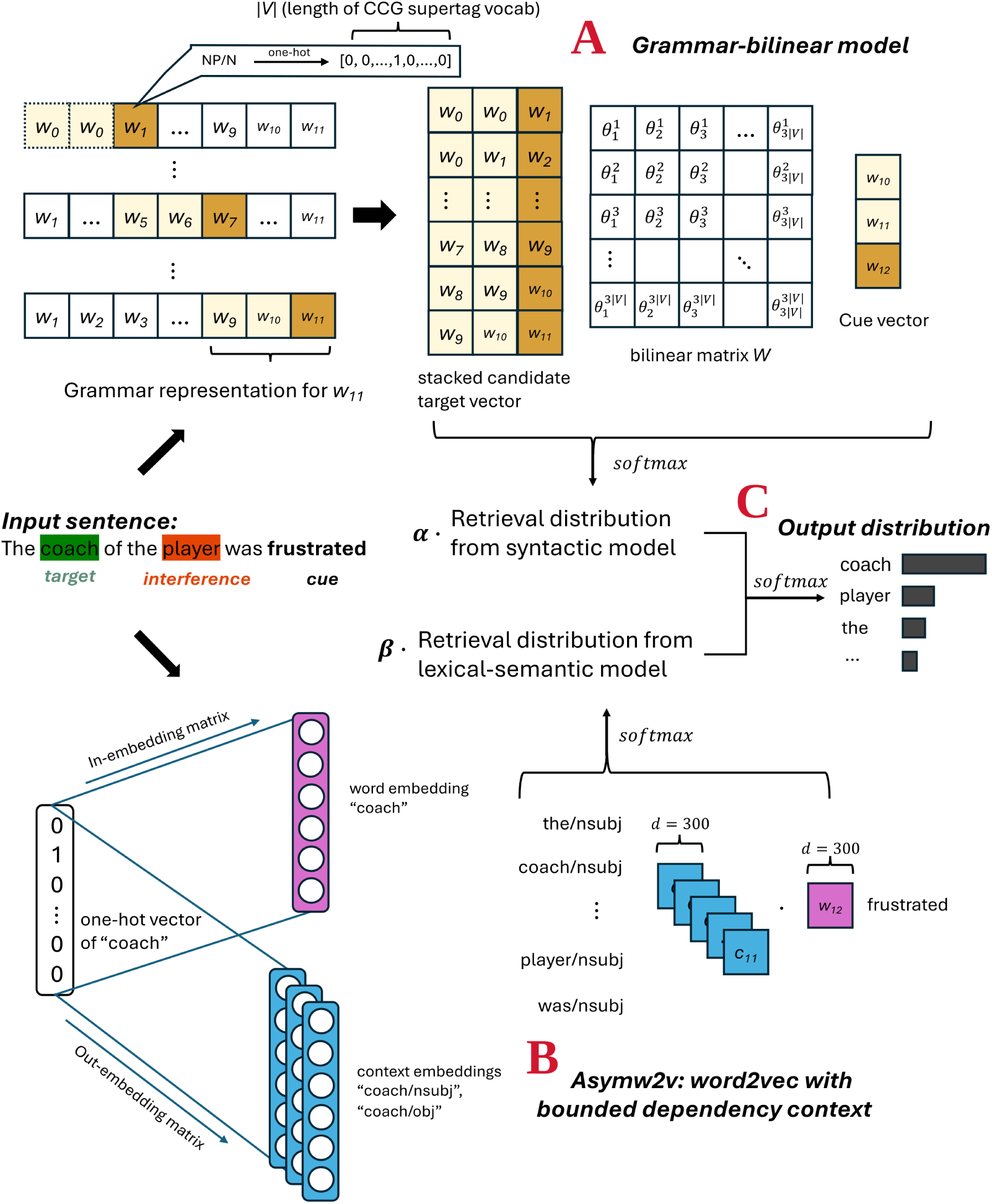
Aspects of the Grammar-Bilinear Asymw2v retrieval model. (A) The grammar-bilinear component represents syntactic context using Combinatory Categorial Grammar supertags and computes cue–target compatibility through a bilinear transformation to produce a syntactic retrieval distribution. (B) The lexical-semantic component uses dependency-based word embeddings to model semantic compatibility between the cue and candidate targets. (C) The two distributions are combined linearly and normalized to produce the final retrieval probability distribution.

#### The Syntactic, Grammar-Bilinear Component

In the syntactic, grammar-bilinear (GB) component (Fig. 4A), syntactic information is approximated using one-hot CCG supertags. CCG is a lexicalist grammatical formalism in which the lexicon assigns each word a richly specified category — a supertag — which encodes its base syntactic type (e.g., noun phrase, sentence/clause), as well as its combinatory potential, including subcategorization and argument structure. Thus, the information encoded in word-by-word supertags is akin to an “almost parse” (Bangalore & Joshi, 1999, p. 237) and provides a rich feature space for modeling fine-grained combinatory constraints in machine learning models (Kasai et al., 2019).

The syntactic environment of word *n*, *h_n_*, is represented as the concatenation of the supertags of that word as well as the two preceding words: *h_n_* = [*G_n−_*_2_; *G_n−_*_1_; *G_n_*], where *G_i_* is a one-hot vector over the vocabulary of supertags. Given this syntactically-informed representation, a learned matrix *W* projects the current word’s representational “cue” into the candidate space, allowing for the calculation of bilinear dot product, softmax-normalized retrieval scores for the words of the preceding context:

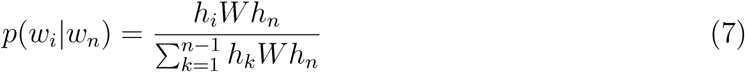

Because supertags encode detailed combinatory constraints, the model learns to assign higher scores to candidates occupying syntactically compatible positions, with structurally incompatible candidates receiving lower retrieval probabilities.

#### The Semantic, Asymmetric Word to Vector Component

The semantic, asymmetric word to vector component (Fig. 4B; Asymw2v) constitutes a dependency-aware Skip-Gram word2vec variant. Vector-embedding models such as word2vec (Mikolov, Sutskever, et al., 2013; Mikolov, Chen, et al., 2013) capture the intuition that words occurring in similar contexts should have similar representations (Firth, 1957; Harris, 1954; Wittgenstein, 1953, see Lenci, 2018 for a computational perspective). In a standard word2vec model, a word’s context is simply the linear window of words surrounding it. The Dependency-Based Word Embeddings model (O. Levy & Goldberg, 2014) used here, however, extends this idea such that context is derived not from surface adjacency, but from syntactic dependency relations — i.e., a word’s context consists of the dependency-labeled words to which it is attached in a dependency graph. This allows for the learning of verb-argument co-occurrence preferences, as well as for the discernment of attribute-level cues such as animacy, in an indirect manner.

In Fig. 1, for example, the word *coach* would have as context not only the adjacent *the* (“*the*/det”), but also the nonadjacent *player* (“*player* /nmod”) and *frustrated* (“*frustrated*/nsubj-1”) — where “-1” indicates that the dependency-labeled context word is the governor in the relation, thus adding even richer detail. As in the Skip-Gram framework, Asymw2v learns two parameter matrices: an input matrix representing words when they serve as “cues,” and an output matrix representing relation-bound contexts. This separation allows the same lexical item to occupy different positions in the embedding space depending on its syntactic role, thus providing a natural way to distinguish circumstances in which a word functions as a retrieval cue versus when it appears as a candidate target within a dependency relation. Retrieval is simulated by using the input embedding for the current word as the cue, and calculating dot product similarity, softmax-normalized retrieval scores using the relation-bound output embeddings for each word in the preceding context.

#### An Interpretable Mixed Model

The full mixed model combines syntactic and lexical retrieval signals via an equally-weighted linear combination of the pre-normalization retrieval scores from the GB and Asymw2vec components, before performing softmax normalization to create the final retrieval probability distribution over the preceding context. This late-fusion formulation reflects cue-based retrieval accounts in which syntactic constraints provide the primary retrieval cues while semantic compatibility contributes graded competition among candidates (Van Dyke & Lewis, 2003; Van Dyke & McElree, 2011). By combining the two signals at the scoring stage, the model preserves strong syntactic filtering while allowing lexical-semantic similarity to modulate retrieval interference.

Having laid out the operating mechanisms of the GBA, we now turn to the sources of the attention patterns analyzed in the current and related work, as well as how these sources relate to cognitive interpretability, or lack thereof. The present GBA model, as applied, is actually an aggregate of individual instances of the mixed GBA model separately trained to predict the correct retrieval target — given an applicable cue word — for many different dependency relations from UD (e.g., nominal subject–nsubj, direct object–obj, case marker–case; see Paragraph GBA NAE). This is in stark contrast to Oh and Schuler (2022) who calculate NAE separately for each of the 12 GPT-2 Small attention heads of the final model layer, before taking the sum across heads — this providing essentially no cognitive interpretability. More grounded, however, is the work of Ryu and Lewis (2021, 2025), who argue that individual GPT-2 attention heads may approximate cue-based retrieval processes through specialization for particular dependency relations — this approach still identifying retrieval-like behavior post hoc within pretrained networks. Another response has been to build smaller models that parameterize retrieval computations more explicitly. Timkey and Linzen (2023), for example, train compact recurrent models with — in keeping with realistic working-memory limits — a single self-attention head. However, because retrieval signals are still implemented through distributed neural dynamics, the mapping between model parameters and specific linguistic constraints remains difficult to characterize mechanistically. In contrast, the GBA model directly parameterizes the retrieval computation, itself.

### Multivariate Temporal Response Function Modeling

One classic way to perform psycho- or neurolinguistic experiments is to present participants with carefully designed stimuli in different conditions and then perform statistical analyses (e.g., ANOVA) to test whether or not the outcome variable (e.g., reading time, fMRI BOLD, EEG µV) differs between conditions. This is not well-compatible with a naturalistic stimulus presentation paradigm, however, where participants simply listen to an audiobook story while their brain data is recorded. While epochs could be constructed for each word and analyzed via single-trial linear regression, this naive approach does not account for (potentially short) temporal spacing between items of interest. For example, a (slightly slow) English spoken word rate of 120 words per minute corresponds to a word onset every 500 ms, on average. If one were to attempt to elucidate the N400 ERP (event-related potential) using some predictor of interest using naturalistic spoken stimuli and single-trial linear regression, they would be forced to face off against variable fluctuations in the target signal attributable just to hearing the next word.

Instead, when using continuous stimuli, observed brain data can be modeled as the linear convolution of some predictor derived from the stimulus with a learned kernel (impulse response), or response function (Brodbeck, Hong, & Simon, 2018; Brodbeck, Presacco, & Simon, 2018; Brodbeck et al., 2020, 2022, 2023; Broderick et al., 2018; Daube et al., 2019; Di Liberto et al., 2015, 2021; Ding & Simon, 2012a, 2012b; Ding et al., 2014; Lalor & Foxe, 2010; Lalor et al., 2006, 2009; Lesenfants et al., 2019; Sohoglu & Davis, 2020, inter alia). Indeed, when multiple predictors are simultaneously considered during response function estimation, the resulting artifact is known as a multivariate temporal response function (mTRF). mTRF analysis allows for the decomposition of a singular brain signal into distinct responses associated with the different predictors entered into the model. In this way, target signal fluctuations which are attributable to aspects of non-interest (e.g., word onset, as in our working example) can be controlled for by adding an associated predictor into the mTRF model.

### Mathematical Formulation

Under the mTRF approach, the target data *y* is modeled as the convolution — i.e., linear filtering — of one or more predictors *x* with a predictor-specific filter kernel, or TRF, *h*. In the case of a single predictor, the value of channel *y* at time point *t* is given by:

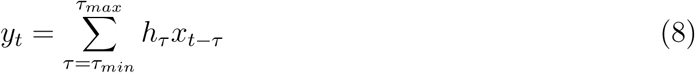

where *x_t_* is the value of predictor time series *x* at time point *t*, *h* is the filter kernel for *x*, and *τ* indexes the temporal lag at which *x* can have an effect on *y*. The extension to an arbitrary number of predictors *n* simply involves the summation of filter responses:

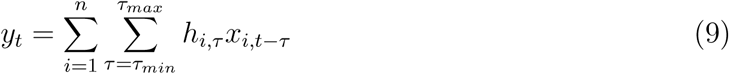

In electrophysiological analysis, *y* and *x* are known, but *h* is not. The encoding or forward modeling task, then, is to estimate the mTRF *h* which best predicts observed brain data from the predictors included in the model. This is algorithmically performed by minimizing the difference between the true brain data *y_t_* and the model-predicted brain data *y*^*_t_*. While there exist other software packages for mTRF estimation — such as the mTRF toolbox for MATLAB (Crosse et al., 2016), which uses ridge regression — we focus on the details of the Eelbrain package for Python (Brodbeck et al., 2023), which uses the boosting algorithm (David and Shamma, 2013; David et al., 2007, see also Friedman et al., 2000; T. Zhang and Yu, 2005). This algorithm is: 1) resistant to overfitting; 2) biased to find sparse mTRF solutions; and 3) is able to handle correlated predictors containing potentially redundant information (David & Shamma, 2013).

### Eelbrain mTRF Estimation via Boosting

First, data are partitioned into separate training and validation splits and an all-zeros mTRF (the *h* term) is created. Then, training data prediction error is measured when each predictor-lag component of the mTRF is independently adjusted both up and down by some small, fixed value, delta. The mTRF is updated to the adjusted mTRF with the greatest error reduction, or, in the case that no improvement was found, delta is either reduced, or training ends if delta has dropped below some prespecified cutoff. After each mTRF update, validation set error is calculated as well, with training ending once validation error stops decreasing.

Because only a single predictor-lag component is adjusted at each step, predictors must compete among themselves in order to explain variance in the observed signal. In this regard, this algorithm can be viewed as coordinate descent, traversing mTRF parameter space in right angles with each update step. As most predictor-lag components stay zero throughout the model estimation process, the algorithm favors sparse solutions. Further, while early stopping based upon validation error serves to prevent overfitting, it additionally promotes sparsity in the final solution by ensuring that no more update steps are taken than are necessary.

While mTRF estimation could be performed once with the entirety of a subject’s available data, in practice — in the tradition of Brodbeck and colleagues (2020, 2022, 2023) — cross-validation is used to calculate “an unbiased estimate of the predictive power of the given predictors” (Brodbeck et al., 2023, p. 22). Under this procedure, each fold, in turn, is used as an independent, held-out test set for the evaluation of an mTRF model estimated on the remaining data. In more detail, for each test fold, each non-test fold is used as the validation fold once, and then the average of these mTRF model estimations is used to make predictions for the held-out test fold. Model fit metrics — i.e., correlation and proportion of variation explained between *y* and *y*^ — are calculated just once, by concatenating the predictions made for the independent, held-out test folds, while the final returned mTRF model is the mean of all of the mTRF models estimated during cross-validation.

### mTRF Analysis Methods

mTRF modeling can be used for computational neurolinguistic analysis in at least two different ways. One technique takes a model comparison approach to identifying whether or not some feature (i.e., predictor) of interest significantly predicts brain activity. The second involves fitting a single, comprehensive model and then analyzing the characteristics of the component TRFs associated with the predictor(s) under consideration.

#### Model Comparison

The predictive power of minimally differing collections of predictors can be used to probe the unique predictivity of some feature of interest, while still controlling for correlated features and features of non-interest. The goodness of an mTRF model — i.e., set of features — can be evaluated by calculating how well its predictions align with the true observed data, as calculated by correlation, *r*, or proportion of variation explained. In this way, hierarchical model comparison can be performed based upon model goodness: a “full” model is estimated with a collection of predictors and then a “reduced” model is estimated containing all of the predictors from the full model, except for the predictor under investigation. If model prediction is better in the full model than in the reduced model, then the conclusion can be drawn that the predictor of interest — present in the full model, but not in the reduced model — contributes to explaining the observed data above and beyond the predictors contained in the diminished model.

#### TRF Analysis

As estimates of the neural response to input impulses of magnitude 1, TRFs can be used to investigate the temporal dynamics associated with the predictors of a model. Thus, they can be thought of as being similar to ERPs or event-related fields (ERFs), but estimated from naturalistic rather than experimental stimuli. Like ERP/Fs, they can similarly be statistically analyzed via mass-univariate tests (e.g., ANOVA to test for a difference among means, *t*-test to test for a difference from 0). They can also be thought of as being similar to the beta coefficient time series resulting from participant-level single-trial linear regression.

### The Present Study

In light of prior work which has correlated disambiguation and memory retrieval complexity metrics derived from broad-coverage neural language models with observed human language processing data, the present project pursues both neural predictivity as well as mechanistic interpretability. To this end, we use source-localized naturalistic MEG data and the mTRF modeling framework. First, a collection of model-goodness comparisons are performed, asking how well do surprisal (derived from the ILCGDP) and NAE (derived from the GBA) perform at modeling observed brain activity, relative to their GPT-2-derived counterparts. Second, taking advantage of the high temporal and moderate spatial resolutions afforded by the MEG neuroimaging modality, we then probe the spatio-temporal dynamics of these predictors when jointly considered within a single comprehensive mTRF model, interpreting the results through the larger scheme of language as a multi-step cognitive process implemented in the human brain.

## Materials and Methods

### MEG Dataset

The MEG dataset used for the present analyses was originally generated as part of the study by Brodbeck et al. (2022). The stimuli comprised 11 excerpts from the audiobook version of *The Botany of Desire*, a non-fiction book by Michael Pollan (2001). Each excerpt had a duration between 210 and 332 seconds, for a total of 46 minutes and 44 seconds. The excerpts were presented in chronological order.

### Participants and Acquisition

Magnetoencephalograms were recorded from 12 native speakers of English (6 female, 6 male; 1 left-handed; mean age = 21 years) with a 157 axial gradiometer KIT MEG system (Kanazawa, Japan) at the University of Maryland, College Park, as they listened to the audiobook excerpts while lying in a supine position. All subjects provided written informed consent in accordance with the University of Maryland Institutional Review Board. Subjects either received course credit (n = 4) or were paid for their participation (n = 8). Each participant answered two to three comprehension questions after each excerpt and was given the opportunity to take a short break. The recordings were made with a sampling rate of 1000 Hz, an online low-pass filter of 200 Hz, and a notch filter at 60 Hz. Further details concerning the data acquisition process can be found in the original paper for this dataset (Brodbeck et al., 2022).

### Preprocessing and Source Localization

Preprocessing of the MEG data was accomplished within the MNE-Python ecosystem (Gramfort et al., 2013, Version 1.9.0) through a near-fully automated pipeline designed to minimize reliance on manual visual inspection steps and to maximize reproducibility. Initial bad channel selection simply followed the bad channels previously identified by Brodbeck et al. (2022). External noise and artifacts were removed through spatiotemporal signal space separation, with a 10 second buffer duration (Taulu & Simola, 2006; Taulu et al., 2004). Data was then filtered 1–40 Hz with a zero-phase FIR filter (MNE-Python default settings).

Independent component analysis (ICA; Bell & Sejnowski, 1995) was then performed with the PICARD algorithm (Ablin et al., 2018) to remove ocular and cardiac artifacts. Following Ferrante et al. (2022), the data were downsampled to 200 Hz prior to ICA for faster convergence. Ocular independent components (ICs) were selected for each participant in a semi-automated manner with mne.preprocessing.corrmap, which used manually selected ICs for one participant (R2614) as a template to select artifactual components for all other participants. Cardiac ICs were automatically detected with the Pearson correlation method. Statistically outlying ICs were then automatically detected with the find_bad_components function from the mne-faster package (van Vliet et al., 2025, Version 1.2.2), a Python implementation of the FASTER artifact rejection pipeline for EEG (Nolan et al., 2010), using the metrics of kurtosis, power gradient, Hurst exponent, and median gradient. Bad channels were interpolated using spherical spline interpolation (Perrin et al., 1989).

Each participant’s digitized head shape was co-registered with the “fsaverage” brain from Freesurfer (Fischl, 2012) using rotation, translation, and uniform scaling. A source space was generated via fourfold icosahedral subdivision of the white matter surface (“ico4”), yielding a total of 2,562 source dipoles per hemisphere. Source dipoles were oriented perpendicularly to the cortical surface. A forward solution was calculated using a single layer boundary element model. Regularized minimum l2 norm current estimates (Dale & Sereno, 1993; Hämäläinen & Ilmoniemi, 1994) were computed for all data using an empty room noise covariance (*λ* = 1/6), yielding signed source time courses which were morphed to fsaverage space.

#### Predictor Variables

Fig. 5 provides a visual summary of the linguistic predictors presented below and elucidates the temporal alignment between brain data, the audio storybook stimulus, acoustic predictors, and word-level predictors.

**Figure 5.**
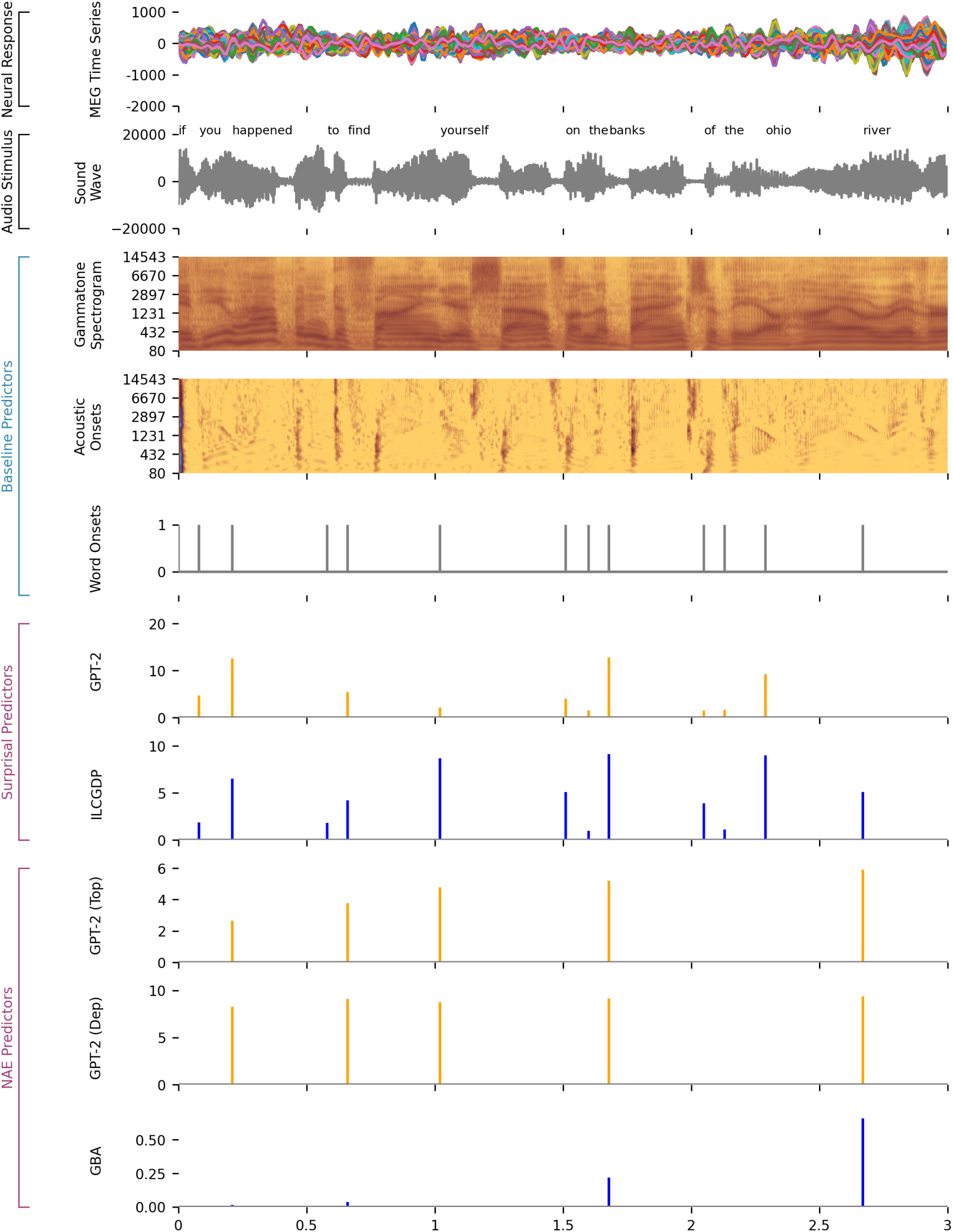
Time series representations of predictors aligned with the MEG data and auditory stimulus. From top: band-pass filtered MEG, raw auditory waveform, gammatone spectrogram, acoustic onset spectrogram, word onsets as uniform impulses, word-wise surprisal according to GPT-2 and the ILCGDP, and word-wise NAE extracted from GPT-2’s topmost layer, GPT-2’s dependency-correlated heads, and the GBA model.

#### Baseline Predictors of Non-Interest

In the modeling of brain data collected during naturalistic language comprehension, it is paramount that low-level predictors of non-interest (viz., acoustics, word rate, lexical frequency) be controlled for.

##### Acoustics

As part of their dataset release, Brodbeck et al. (2022) include acoustic features extracted from the audiobook stimulus. Eight features were extracted from an eight band gammatone spectrogram, and eight features were extracted from an eight band acoustic onset spectrogram, “both covering frequencies from 20 to 5000 Hz in equivalent rectangular bandwidth space (Heeris, 2018) and scaled with exponent 0.6 (Biesmans et al., 2017)” (Brodbeck et al., 2022, p. 19). Gammatone filters model peripheral auditory system response characteristics, and acoustic onset spectrogram calculations model acoustic onset detection (Fishbach et al., 2001).

##### Word Rate and Annotating Word-Level Predictors

Brodbeck et al. (2022) also include Praat TextGrids (Boersma & Weenink, 2026) for the audiobook, which were generated by combining the Carnegie Mellon University pronunciation dictionary with the Montreal Forced Aligner (McAuliffe et al., 2017). Following Brodbeck et al. (2023), word-level predictors are annotated at word-onset. Word rate is a particular instantiation which marks word onset with value 1 and which is meant to account for variance attributable just to hearing words.

##### Lexical Frequency

The log-transformed frequency of each word in the audiobook was calculated relative to a subset of the Open American National Corpus (American National Corpus Project, 2015). The sub-corpus contained 4,491,322 tokens across 61,341 documents. Log frequency was calculated by taking the logarithm of each token’s frequency count. Following Brodbeck (2024), we further imposed an inverse transform on the log frequency values by computing 17 minus log frequency, creating an inverse frequency measure where higher values correspond to less frequent words.

#### Predictors of Interest

We now turn our attention to the predictors which are of interest to the current project: 1) surprisal — our operationalization of disambiguation — derived from the ILCGDP and GPT-2; and 2) normalized attention entropy — our operationalization of memory retrieval — derived from the GBA and GPT-2, as well.

##### ILCGDP Surprisal

Audiobook sentences were parsed with the ILCGDP using basic Stanford Dependencies (de Marneffe & Manning, 2008a, 2008b; de Marneffe et al., 2006), a beam size of 64, and part of speech (POS) tags predicted by spaCy (Honnibal et al., 2020). Because the ILCGDP does not include POS tag in the joint probability distribution — i.e., it does not predict POS tags (cf. Buys & Blunsom, 2015a, 2015b) — it is simply a case of a natural language processing pipeline to utilize the POS tags predicted by spaCy as features. Word-by-word surprisal was calculated as the log ratio of the prefix probabilities before and after a word, with the first word of a each sentence being assigned a surprisal value of 0.

##### GPT-2 Surprisal

Following the finding by Oh and Schuler (2023) and Oh et al. (2022) that the efficacy of a large Transformer LM to predict reading times decreases as model size and quality increases, we follow the trend in the recent computational psycho-and neurolinguistic literature (Goldstein et al., 2022; Hu et al., 2020; Huang et al., 2024; Wilcox et al., 2020, inter alia) of calculating surprisal from GPT-2 Small (Radford et al., 2019). We calculated GPT-2 surprisal values with code using Hugging Face Transformers (Wolf et al., 2020). First, log softmax was applied over output logits to derive token-wise log probabilities. These were then used to compute token-wise surprisals by taking their negative logarithm. Following the approach of Ryu (2025), whenever GPT-2’s byte-pair encoding tokenizer recognized a word as a combination of multiple sub-lexical tokens, we took the maximum surprisal of the sub-lexical tokens as representative of the surprisal of the word. The first word of each sentence was assigned a surprisal value of 0.

##### GBA NAE

In the current implementation, mixed GBA models were trained to implement cue-based memory retrieval for the 23 UD dependency relations listed in Table 1. The selected relations are of the sort that have been used in previous computational neurolinguistic work (Kumar et al., 2024; Lopopolo et al., 2021). For each relation, both governors and governees were allowed to serve as retrieval cues, targeting their grammatical counterpart if it occurred in the prior context of the sentence. For example, under the nsubj relation, a verb may retrieve its subject noun from prior context, while in constructions such as passives, a subject noun may retrieve its governing verb if that verb precedes it. Each GBA instance is therefore relation-specific: it assigns retrieval scores only to tokens participating in the dependency it was trained to model. It should be noted that, unlike GPT-2, which distributes attention over all prior tokens regardless of grammatical type, the GBA model does not attempt to model retrieval operations beyond the scope of the grammatical relation it is trained for. The GBA nsubj instance, then, is only able to assign an NAE value to verbs and nouns. On all other tokens, it assigns an unconsidered NA value.

**Table 1.** The 23 Universal Dependencies relations considered by the aggregate mixed Grammar-Bilinear Asymw2v model (see Fig. 4).

| Abbreviation | Full Name |
| --- | --- |
| acl | adnominal clause |
| acl:relcl | relative clause |
| advmod | adverbial modifier |
| amod | adjectival modifier |
| appos | appositional modifier |
| aux | auxiliary |
| case | case marker |
| cc | coordination conjunction |
| ccomp | closed clausal complement |
| compound | compound |
| conj | conjunction |
| cop | copula |
| det | determiner |
| discourse | discourse element |
| iobj | indirect object |
| mark | marker |
| nmod | nominal modifier |
| nsubj | nominal subject |
| nummod | numeral modifier |
| obj | object |
| obl | oblique |
| parataxis | parataxis |
| xcomp | open clausal complement |

Aggregating across GBA instances for all 23 dependencies, results in the annotation of approximately 37.6% of the audiobook with NAE values. This is close to the theoretical maximum coverage that is achievable with the constraint that retrievals only go leftward (3,053 possible cue tokens, 7,743 total tokens, theoretical maximum coverage = 39.4%).

Aggregation was simply implemented as summing across the NAE values assigned by each GBA instance to a word.

##### GPT-2 NAE

As introduced, there exist multiple possible ways to derive NAE values from GPT-2, each operationalizing a different Transformer LM-based hypothesis. In this study, we are concerned with two such approaches, both involving the aggregation of NAE values from a set of GPT-2 Small attention heads, with each head analogous to a GBA instance as described above. Likewise, then, aggregation is implemented as simply summing across the NAE values calculated by the attention heads.

The first approach, following Oh and Schuler (2022), is to hypothesize that the attention heads most relevant to memory retrieval in human sentence processing are those of the topmost layer (layer 12), since this is the layer which directly informs GPT-2’s next-token predictions. Under this account, the retrieval operations implemented by the 12 heads of layer 12 are sufficient to explain cue-based memory retrieval in humans. It remains uninterpreted, however, what the targets being retrieved in the operations conducted by these heads are.

The second approach is to aggregate across only those attention heads whose attention patterns have been demonstrated to correlate with expected retrieval operations for specific grammatical relations. Following a method proposed by Voita et al. (2019), Ryu (2025) identified GPT-2 heads across different layers whose attention patterns correlated with retrieval operations along particular grammatical relations. This implements the hypothesis that only some attention heads contribute meaningfully to memory retrieval whereas irrelevant heads contribute noise (Ryu, 2025, p. 22). It remains unclear, though, what is being retrieved when a grammatically-specialized head encounters a token for which its grammatical specialty is irrelevant.

These two approaches bear out two different configurations of GPT-2 Small, and we refer to the NAE values extracted from each configuration as GPT-2_Top_NAE and GPT-2_Dep_NAE respectively. With regard to annotation coverage, for fairness of comparison, we zero-masked both GPT-2_Top_NAE and GPT-2_Dep_NAE to the same 37.6% of words annotated by the GBA implementation.

### mTRF Model Estimation

In the present project, our goals are twofold. First, we seek to evaluate whether surprisal and NAE derived from our interpretable models — the ILCGDP and the GBA — are as equally adept at explaining naturalistic MEG source space data as their counterparts from GPT-2. In other words, are our bespoke algorithmic models at least as aligned with the brain as GPT-2 is during naturalistic listening? Second, we wish to elucidate the spatio-temporal dynamics of surprisal and NAE effects in sentence comprehension when they are jointly considered. That is, are the effects of ILCGDP surprisal and those of GBA NAE dissociable in space and/or time? To these two ends, we estimated a number of mTRF models for our analyses.

#### Model Estimation

mTRF models were estimated using the TRF-Tools extension (version 11; Brodbeck and Hyde, 2025) for the Eelbrain package (version 0.41; Brodbeck, 2025; Brodbeck et al., 2023). For mTRF estimation, the source-localized MEG data were downsampled to 50 Hz and the 11 separate excerpt sections were concatenated together. mTRFs were computed independently for each subject and each virtual source dipole (full ico4 source space) using 50 ms wide Hamming window basis functions at time lags from *τ_min_* = −100 ms to *τ_max_* = 1000 ms. All responses and predictors were standardized by centering and dividing by the standard deviation. TRF kernels were estimated using the boosting algorithm in a fourfold cross validation regimen, with the training objective of minimizing L2 error. Selective stopping was employed — whenever a training step caused an increase in error in the validation segment, the TRF for the specific predictor responsible for the increase was frozen, while training continued for the remaining predictors. Training continued until all individual predictor TRFs were frozen.

Seven separate mTRF models were fitted for the following seven sets of predictors:

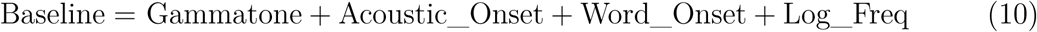

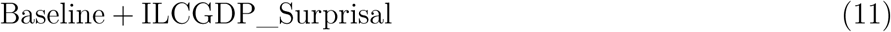

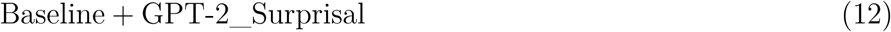

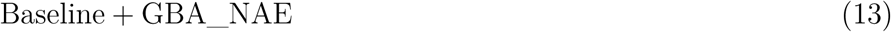

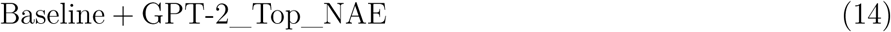

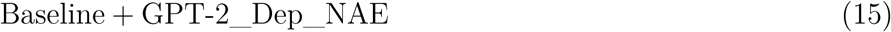

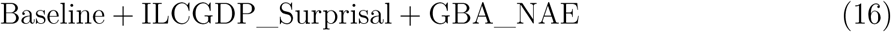

where Gammatone and Acoustic_Onset indicate the eight-band gammatone spectrogram and eight-band acoustic onset spectrogram predictors from Brodbeck et al. (2022), respectively; GPT-2_Top_NAE indicates GPT-2 NAE derived from the topmost Transformer layer; and GPT-2_Dep_NAE indicates GPT-2 NAE derived from the grammatically-specialized attention heads described in Ryu (2025). Model comparison analyses relied on formulae 10–15, while the model specified in formula 16 was used in the spatio-temporal TRF analysis.

### Model Comparison Analyses

#### First Level (Within-Subject Analysis)

We quantified the goodness-of-fit of an mTRF model — i.e., its predictive power over the brain data — as the proportion of variation explained (PVE) for the true response by the predicted response. To isolate a specific predictor of interest’s unique contribution to PVE, we take a “reduced” baseline model (formula 10; containing the baseline predictors of non-interest) and a minimally different “full” model containing baseline predictors plus the predictor of interest, and then compute the gain in PVE that results from including the predictor of interest (following Crabbé et al., 2019; Franzluebbers et al., 2024; Wolfman et al., 2024). This is done per source-space vertex and collapses over the time dimension — yielding a scalar PVE gain value at each vertex. For each participant, we computed how much each predictor of interest (model formulae 11–15) increased PVE over the baseline model.

Next, the PVE gain maps for the predictors derived from GPT-2 were subtracted from the PVE gain maps derived from the ILCGDP and the GBA, setting up 3 contrasts:

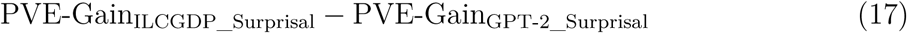

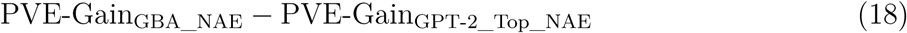

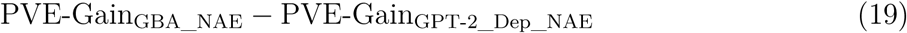

Under this setup: a positive value (at a vertex) would indicate that the predictor derived from the interpretable model is more well-aligned with the observed brain data than the Transformer model (GPT-2); a value of 0 would indicate that both models are equally brain-aligned; and a negative value would indicate that the Transformer model is more aligned with the brain than the cognitive model. Thus, at the first level, each participant contributes 3 contrast brain maps, where each map is a high-dimensional data point that tells us how much more a interpretable predictor increases PVE compared with a GPT-2 predictor.

#### Second Level (Group Analysis) and Statistical Evaluation

At the second (group) level, using Eelbrain, these contrast maps were submitted to two-tailed, mass-univariate, one-sample *t*-tests to find source-space clusters where the contrast between the interpretable predictor and the GPT-2 predictor is statistically significant. First, *t*-tests were computed at each vertex to determine whether their contrast value was significantly different from a population mean of 0 across participants. This produces a *t*-map of *t*-values at each vertex (along with corresponding *p*-values). To estimate significance at the group level, spatial cluster-based permutation testing was performed over the resulting *t*-maps (Maris & Oostenveld, 2007). This non-parametric method first identifies clusters of significant responses that are contiguous in space, then bootstraps a null distribution from the data to estimate cluster statistics. Adjacent vertices (of the same sign) were clustered together if their uncorrected *p*-value was *p <* 0.05.^4^ The *t*-values within each cluster were summed to calculate a cluster-level statistic. A null cluster distribution was estimated by repeating this procedure for 4,095 permutations^5^ in which the spatial values of each participant were randomly shuffled with 0, simulating data under the null hypothesis. Cluster *p*-values were determined by the position in the null distribution, with clusters in the bottom or top 2.5% considered statistically significant. This method accounts for multiple (within-analysis) comparisons (through the permutation-based null distribution) while acknowledging the non-independence of adjacent vertices in source space.

### Analysis Regions

While mTRF models were fit to all vertices in the ico4 source space, statistical analysis was limited to the dorsal and ventral streams of the bilateral language network (Bornkessel-Schlesewsky & Schlesewsky, 2013; Friederici & Gierhan, 2013; Hickok & Poeppel, 2007; Matchin & Hickok, 2020). Using the “aparc” parcellation scheme in Freesurfer, otherwise known as the Desikan-Killiany (DK) atlas (Desikan et al., 2006), the dorsal stream was defined to include the inferior frontal gyrus (IFG; DK parcels: pars opercularis, pars orbitalis, pars triangularis), the posterior superior temporal gyrus (pSTG; DK parcels: superior temporal gyrus, banks of superior temporal sulcus, transverse temporal gyrus), and the inferior parietal lobule (DK parcels: inferior parietal lobe, supramarginal gyrus). The ventral stream comprised the same IFG and pSTG regions plus the anterior temporal lobe (DK parcels: temporal pole, middle temporal gyrus, inferior temporal gyrus).

### Spatio-Temporal TRF Analysis

To investigate the spatio-temporal dynamics of ILCGDP surprisal and GBA NAE, we statistically analyzed the TRFs estimated by the large mTRF model specified in formula 16. At the group level, the surprisal and NAE TRFs (analogous to subject-level ERP/Fs or single-trial linear regression beta coefficient time series) were submitted to two-tailed, mass-univariate, one-sample *t*-tests to determine, at each vertex and time point, whether their values are significantly different from a population mean of 0 across participants. This produces a *t*-value at each vertex and time point. While the mTRF models were estimated across a time window spanning −100 to 1000 ms, we constrain our TRF analysis to the 200–1000 ms window, since high-level language-related cognitive processes are unlikely to be associated with true effects at earlier latencies.

For estimating significance at the group level, spatio-temporal cluster-based permutation testing was then performed over the resulting vertex × time *t*-maps. Clusters of significant responses that were contiguous in space and time were first identified before bootstrapping a null distribution from the data to estimate cluster-level statistics. Adjacent time points and vertices (of the same sign) were clustered together if their uncorrected *p*-value was *p <* 0.05. *T*-values were summed to calculate a cluster-level statistic. A null cluster distribution was estimated by repeating this procedure for 4,095 permutations in which the spatio-temporal values of the TRF of each participant were randomly shuffled with 0, simulating data under the null hypothesis. Cluster *p*-values were determined by the position in the null distribution, with clusters in the bottom or top 2.5% considered statistically significant. Like the model comparison analyses, the TRF analysis was constrained to vertices falling within the bilateral language network.

### Data and Code Availability

The MEG data and predictors provided by Brodbeck et al. (2022) are available at https://datadryad.org/dataset/doi:10.5061/dryad.nvx0k6dv0. The remaining predictors, model estimation code, and statistical analysis code are available at https://osf.io/prv45/overview?view_only=ea1da7be5a2142ae9fff43ee667a46f4.

## Results

We present our results with uncorrected, cluster-level *p*-values. However, to account for multiple analyses of the same underlying data, we apply a conservative Bonferroni correction for the interpretation of significance. Specifically, we perform three analyses to compare increases in the proportion of variation explained — once for surprisal and twice for normalized attention entropy. Additionally, we analyze the TRFs associated with ILCGDP surprisal and GBA NAE in a large singular model. To correct for these multiple analyses, we consider cluster-level results significant only at a corrected threshold of *α* = 0.05 / 4, which corresponds to an uncorrected *p* < 0.0125.

### Model Comparison Analyses

Model comparisons revealed that complexity metrics derived from the ILCGDP and GBA models presented here are no less well-aligned with the brain than their counterparts derived from the Transformer model GPT-2 Small. Specifically, we tested whether PVE gains induced by our interpretable model predictors differed significantly from those induced by GPT-2-derived predictors across language network source space. All clusters failed to reach significance, not only at the corrected threshold, but even at a more lax uncorrected *p <* 0.05 threshold. This indicates that, in no cortical language network region, does ILCGDP surprisal predict brain activity any worse than GPT-2 surprisal, nor does GBA NAE predict brain activity any worse than either of the GPT-2-derived NAE predictors.

### Spatio-Temporal TRF Analysis

A number of ILCGDP surprisal and GBA NAE TRF clusters reached the corrected threshold of *p <* 0.0125. These are shown in Figs. 6 and 7, respectively.

**Figure 6.**
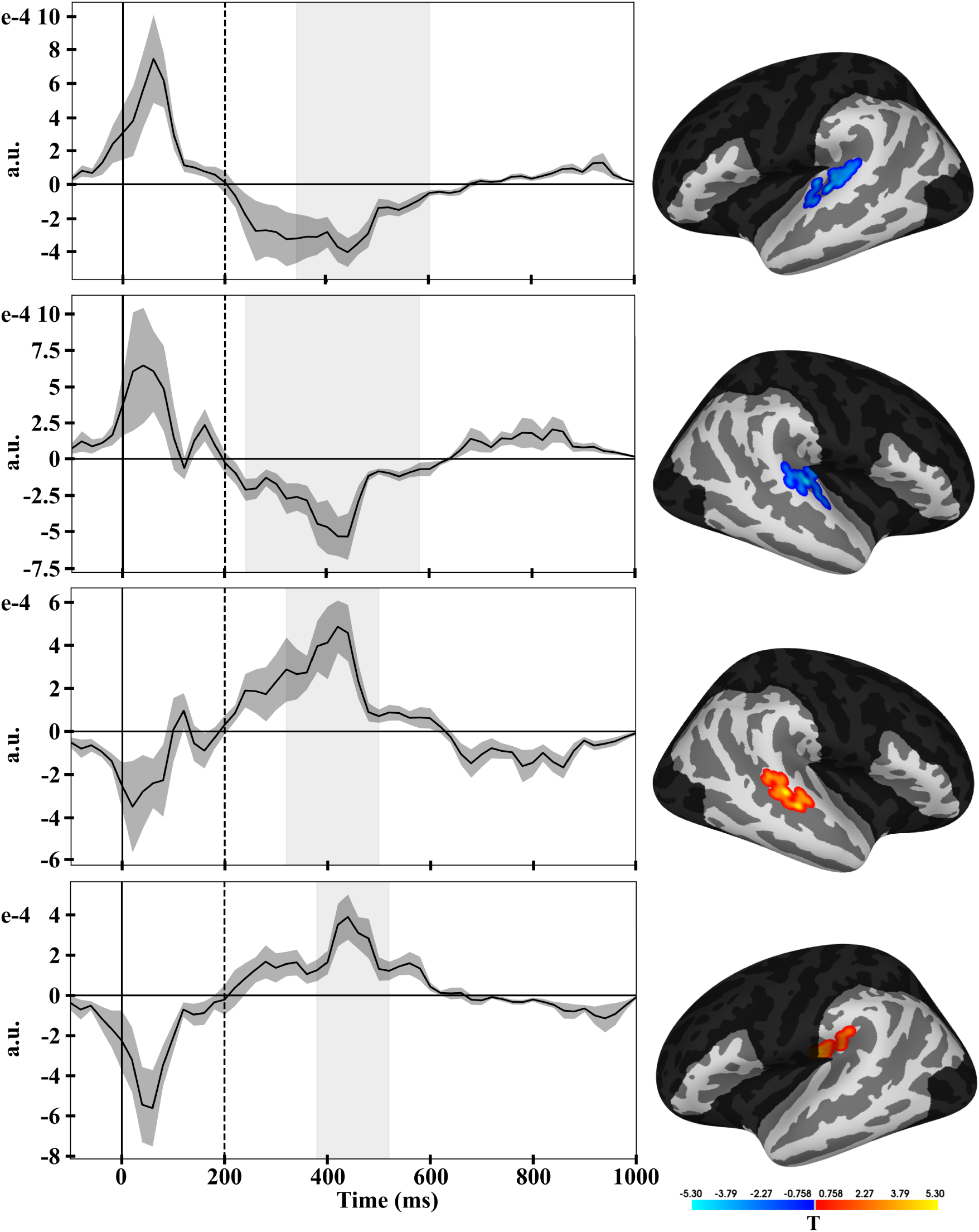
Incremental Left-Corner Generative Dependency Parser surprisal clusters. Brains show the distribution of t-values and vertices included in each cluster at cluster peak, with the spatial extent of the analyzed bilateral language network demarcated. Time series show the average temporal response function in these vertices, with the temporal extent of the cluster shaded. Dashed line demarcates the 200–1000 ms temporal region of interest. a.u.: arbitrary units.

**Figure 7.**
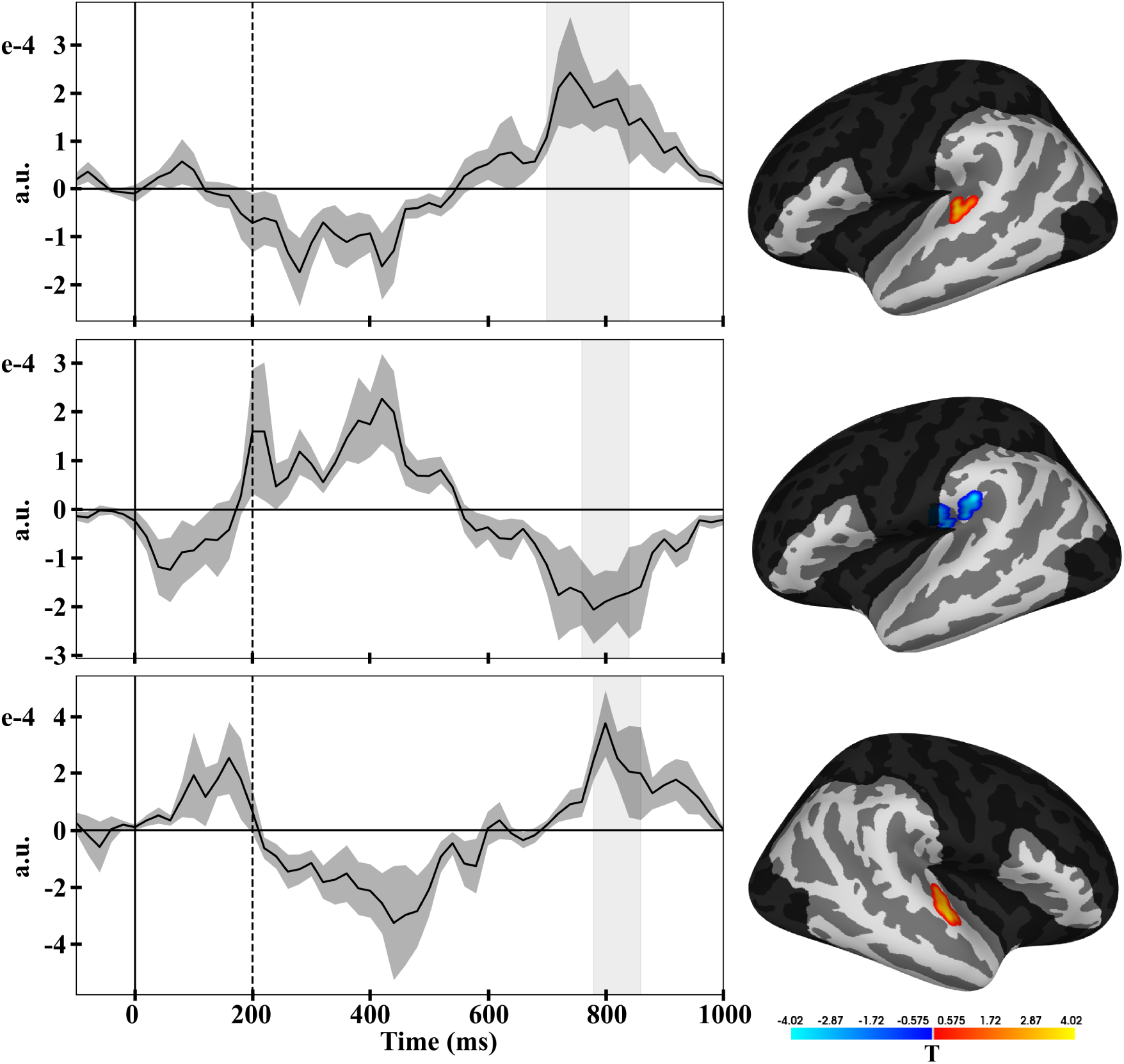
Grammar Bilinear Asymw2v normalized attention entropy clusters. Brains show the distribution of t-values and vertices included in each cluster at cluster peak, with the spatial extent of the analyzed bilateral language network demarcated. Time series show the average temporal response function in these vertices, with the temporal extent of the cluster shaded. Dashed line demarcates the 200–1000 ms temporal region of interest. a.u.: arbitrary units.

#### ILCGDP Surprisal

Four clusters were identified for surprisal derived from the ILCGDP: 1) a negative-correlation cluster consisting of 37 vertices in the left superior temporal gyrus (STG) 340–600 ms (*p <* 0.001); 2) a negative-correlation cluster consisting of 27 vertices in the right STG 240–580 ms (*p <* 0.001); 3) a positive-correlation cluster consisting of 24 vertices in the right STG 320–500 ms (*p* = 0.003); and 4) a positive-correlation cluster consisting of 16 vertices in the left supramarginal gyrus (SMG) 380–520 ms (*p* = 0.009).

#### GBA NAE

Three clusters were identified for GBA NAE: 1) a positive-correlation cluster consisting of 22 vertices in the left STG 700–840 ms (*p <* 0.001); 2) a negative-correlation cluster consisting of 18 vertices in the left SMG 760–840 ms (*p <* 0.001); and 3) a positive-correlation cluster consisting of 8 vertices in the right STG 780–860 ms (*p* = 0.005).

#### Early Responses

Noticeable in both Fig. 6 as well as Fig. 7 are early TRF responses in the 0–200 ms time window which was excluded from group-level statistical analysis. While the boosting algorithm for mTRF estimation (David & Shamma, 2013; David et al., 2007) has been argued to be robust to correlated predictors containing potentially redundant information and biased toward sparse (i.e., flat, zero-valued) mTRF solutions (Brodbeck et al., 2023), we attribute these peaks to “contamination” resulting from predictor correlation between surprisal and NAE with word rate (the response to simply encountering a word onset). While it was not known a priori that this would be observed, the 0–200 ms time window was excluded beforehand from group-level statistical analysis on the assumption that true word-conditioned effects from cognitive operations like disambiguation and memory retrieval should not occur that early.

#### TRF Signedness

As introduced, the mTRF models were fitted to signed source space data following the recommendation of Brodbeck, Presacco, and Simon (2018, p. 165): “when analyzing continuous responses, where high pass filtering replaces baseline correction, a change of the sign in the current estimate is important information. Hence, directional (‘fixed orientation’) source estimates, which preserve information about the dipole orientation, are preferable.” This means that spatio-temporal cluster interpretation requires care because anatomical features “lead to source estimates with alternating sign across gyri and sulci due to alternating cortical surface orientation” (ibid., p. 166). To illustrate the point, it is likely the case that the two right hemisphere surprisal clusters shown in Fig. 6 belong to the same underlying brain response, but owing to perpendicular, fixed dipole orientation, the sign of the minimum norm current estimate switches between the superior temporal gyrus and the superior temporal sulcus, thus leading to two temporally overlapping and spatially adjacent clusters with opposite sign (see also Brodbeck, Presacco, and Simon, 2018 Fig. 3 and surrounding discussion). For this reason, the discussion section emphasizes spatio-temporal TRF cluster timing more so than location (moderate emphasis), and directionality (no emphasis).

## Discussion

By reaching “for the goal of scientific explanation using computational models that are by and large interpretable,” the present project has taken up the neurocomputational modeling call to action put forth by Hale et al. (2022, p. 9). We first remark on the demonstration that — despite the prominence of neural large language models (LLMs) in the contemporary psycho- and neurolinguistic literature — smaller models containing human-grounded inductive biases can be just as well aligned with language network brain activity recorded during narrative listening. In other words, the computational cognitive models presented here achieve equal ability as LLMs, with the benefits of both interpretability as well as data and compute efficiency.

Following this, we discuss a staged processing (Sternberg, 1969) account of word-by-word sentence processing based upon our results correlating ILCGDP surprisal and GBA NAE with naturalstic MEG data. While the idea that incremental sentence comprehension proceeds as an ordered collection of operations is not a novel one (see Bornkessel & Schlesewsky, 2006; Frazier & Fodor, 1978; Friederici, 2002), the present project does, however, demonstrate the applicability of such lines of reasoning at an unprecedented scale — owing to the broad-coverage statistical nature of the computational models used, as well as the naturalness of the analyzed data.

### Equivalence in Predictive Power

Within computational cognitive neuroscience, models of cognition are often judged by the merits of their predictive power — how well they are able to predict neural data — quantified by “brain score” metrics such as correlation or proportion of variation explained. Yet, predictive power does not an explanatory model make. As emphasized by Shmueli (2011), predictive modeling and explanatory modeling are different (but mutually supportive) activities. Prediction is concerned with capturing statistical associations while explanation involves inferring causal relations. Notably, neither necessarily entails the other. Here, we adopt the ideal that a good scientific model should be both predictive as well as explanatory — that is, the model should provide an interpretable mapping between the theory that it implements as well as the predictions that it makes (Guest & Martin, 2021). On this point, the mechanistic interpretability of a given neurocomputational model, along with its alignment with brain data, are both paramount, neither sufficient on their own (Hale et al., 2022).

This is not to say that LLMs lack any explanatory power, however. For example, recent work by Ryu and Lewis (2021, 2025) proposes that specific attention heads in GPT-2 implement cue-based memory retrieval for specific grammatical dependency relations (e.g., a verb retrieving its subject). Yet, gaps in interpretability do remain. For example, it is unclear what such a grammar-specialized attention head is attempting to retrieve when the cue token is unrelated to its specialty. In other words, the GPT-2 theory underspecifies what features are targeted in retrieval operations. This is not to downplay the progress made in regard to “unblackboxing” GPT-2. Nevertheless, it must be acknowledged that GPT-2 has yet to yield an adequately specified explanatory account of the human sentence processor. In contrast, the ILCGDP and GBA are fully interpretable models. As a scientific principle, between any two models with equal predictive power but unequal explanatory power, it should be concluded that the model with superior explanatory power is to be preferred.

### A Staged Theory of Sentence Processing

While there has been much work theorizing and investigating disambiguation and memory retrieval as separate accounts, less work has considered both of these aspects of language comprehension jointly (although see Campanelli et al., 2018; Tung & Brennan, 2023). In temporally dissociating surprisal and NAE, we build off of recent work by Ryu and Lewis (2025), and advocate for a staged theory of sentence processing. Without excluding other potentially relevant dimensions, and focusing just on those investigated here, one stage — indexed by surprisal — reflects early lexical disambiguation, while another later stage — indexed by normalized attention entropy — reflects memory retrieval and integration, i.e., dependency construction.

#### Surprisal

A number of spatio-temporal clusters were observed for ILCGDP surprisal in the bilateral STG and left SMG ∼300–550 ms. While the N400 is technically an EEG ERP, the results of previous research lead to an interpretation of the present collection of MEG effects as homologous. The N400 is a negative-going, centro-parietal ERP, peaking approximately 400 ms following stimulus onset. While once thought to be found for words which are semantically anomalous, given a context (Kutas & Hillyard, 1980; Osterhout & Holcomb, 1992; Osterhout & Mobley, 1995), the N400 is now thought to be observed for all content words and to reflect inverse Cloze probability (Phillips et al., 2005). Regarding previous staged processing proposals, the N400 fall under Phase 2 of the model presented in Friederici (2002) — which involves lexical-semantic and morphosyntactic processing for thematic role assignment — and Phase 2 in the Extended Argument Dependency Model (eADM) of Bornkessel and Schlesewsky (2006) — which involves the establishment of both inter-argument as well as argument-verb relations.

Previous computational modeling work has associated neural network language model-derived surprisal with the N400 both in EEG (Chen et al., 2025; Frank et al., 2013, 2015; Heilbron et al., 2022) and in MEG (Heilbron et al., 2022), thus positively linking surprisal with the processing difficulty of a word, given context. Other EEG results have had less N400-typical spatial distributions, while remaining within the temporal range identified here. Hale et al. (2018), for example, find that surprisal derived from Recurrent Neural Network Grammar (Dyer et al., 2016) is associated with an early frontal positivity ∼250 ms, while Brennan and Hale (2019) find that probabilistic context-free grammar-derived surprisal is associated with an anterior negativity (216–554 ms) for content words, and an anterior positivity (210–310 ms) for grammatical function words.

From the temporal domain, now, to the spatial domain, it has been shown that language network (e.g., Matchin & Hickok, 2020) activation, broadly, correlates with surprisal (Oh et al., 2021, 2022; Shain et al., 2020). More specifically, however, the clusters identified here are in alignment with results which have associated surprisal with activation in left posterior temporal (Brennan et al., 2016, 2020; Willems et al., 2016), left inferior parietal (Brennan et al., 2016, 2020; Lopopolo et al., 2017; Willems et al., 2016), and bilateral superior temporal (Franzluebbers et al., 2024; Lopopolo et al., 2017; Stanojević et al., 2023; Wolfman et al., 2024) cortex.

Two additional interpretations are worth briefly considering. The first is that the observed surprisal clusters are correlating not with the N400, but with the left anterior negativity (LAN; Coulson et al., 1998; Hoen & Dominey, 2000; Kaan & Swaab, 2003a; Kluender & Kutas, 1993; Molinaro et al., 2011; Neville et al., 1991; Osterhout & Mobley, 1995, inter alia), an ERP which is associated with morphoysyntactic expectance violation, and which also falls under Phase 2 of both the Friederici (2002) model and the eADM (Bornkessel & Schlesewsky, 2006). While this cannot be ruled out, it seems less likely, as previous work investigating the ability of surprisal to predict various ERP components shows reliable effects for the N400 only, and not for the LAN (Frank et al., 2015). The second is that surprisal is indexing the M350 ERF (Embick et al., 2001; Pylkkänen & Marantz, 2003; Pylkkänen et al., 2002, 2004; Solomyak & Marantz, 2010; Stockall et al., 2004; Zipse et al., 2006, inter alia), a component which is associated with lexical processing and which has been investigated using properties such as lexical frequency, neighborhood density, and phonotactic probability. The commonly applied single-item presentation paradigm, however, is a far cry from the continuous, naturalistic speech stimulus used here. While is has been shown that M350 latency correlates with lexical frequency (Embick et al., 2001), this is included as a predictor of non-interest in the large mTRF model specified in formula 16. Taken together, it seems unlikely that surprisal is correlating with this component.

#### Normalized Attention Entropy

More interesting are the spatio-temporal clusters observed for GBA NAE in the bilateral STG and left SMG ∼750–850 ms. While deviations from zero were also observed in these clusters ∼300–500 ms (Fig. 7), no clusters falling within this early time window reached significance, even at a lax uncorrected *p <* 0.05. The only other work investigate NAE in the context of neuroimaging is that of Terpstra et al. (2025), who find that NAE is predictive of N400 amplitude in naturalistic reading above and beyond surprisal. It should be noted, however, that their analysis investigates only N400 amplitude, i.e., they do not entertain later time windows, like we do here.

Once again, while the P600 is technically an EEG ERP, the results of previous research (expounded upon below) lead to an interpretation of the present collection of MEG effects as homologous. The P600 is a positive-going, centro-parietal ERP which begins approximately 500 ms after the presentation of the stimulus and is thought to index retrieval and/or integration difficulty (Aurnhammer et al., 2021; Fiebach et al., 2001, 2002; Kaan & Swaab, 2003b; Kaan et al., 2000; Molinaro et al., 2011; Phillips et al., 2005). In terms of previous staged processing accounts, the P600 corresponds to Phase 3 of the model presented in Friederici (2002) — which involves information integration — and Phase 2 in the eADM (Bornkessel & Schlesewsky, 2006) — i.e., argument-argument and argument-verb relation establishment.

With regard to localization, strong evidence for involvement of the STG in cue-based memory retrieval comes from Glaser et al. (2013), who adapt the 2 x 2 high vs. low, semantic vs. syntactic interference during subject retrieval experiment of Van Dyke (2007) to fMRI. They find increased activation in left STG for both high compared to low, syntactic as well as semantic interference. Less direct support comes from implications of the posterior temporal lobe in antecedent retrieval (Hammer et al., 2011; S. Zhang et al., 2022) and work which casts memory demand in the maintenance more so than retrieval light (Fiebach et al., 2001, 2005, 2007).

Next, we present a number of carefully developed associations to relate: 1) normalized attention entropy; 2) similarity-based interference during cue-based memory retrieval; and 3) late electrophysiological effects. Interference occurs during memory retrieval when multiple elements in memory match, to some degree, the retrieval cues — with the non-target interfering elements known as distractors (see Vasishth & Engelmann, 2021, for a book-length treatment). The psycholinguistic consensus is that ungrammatical sentences like (2a) are easier to process than those like (2b; Dillon et al., 2013; Jäger et al., 2017, 2020; Lago et al., 2015; Pearlmutter et al., 1999; Tanner et al., 2014, 2017; Tucker et al., 2015; Wagers et al., 2009, inter alia; examples adapted from Bock and Miller, 1991, 1. p. 45).

(2) a. *The cost 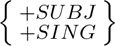 of the improvements 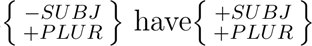 not yet been estimated.

b. *The cost 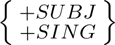 of the improvement 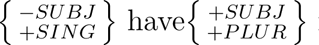 not yet been estimated.

This phenomenon — also known as agreement attraction^6^ — is thought to result from facilitatory interference, i.e., a reduction in the cost of processing an ungrammatical sentence due to the presence of an interfering distractor. In both (2a) as well as (2b), at the verb *have*, a retrieval is made for the subject of the sentence using the cues {+*SUBJ*} and {+*PLUR*}. In both, *have* will partially match the true subject *cost* on {+*SUBJ*}. Only in (2a), however, will *have* additionally partially match *improvements* on {+*PLUR*}, thus leading to an amelioration of the effect of ungrammaticality. Further consideration of the possible cognitive-psychological details causing facilitatory interference in ungrammatical stimuli are beyond the scope of the present project, however the reader is referred to Dillon et al. (2013), Wagers et al. (2009), and Vasishth and Engelmann (2021).

Having introduced the the phenomenon of facilitatory interference during the processing of ungrammatical agreement attraction constructions, we now associate self-attention diffuseness with late electrophysiological effects. First, EEG results show that P600 effects elicited for ungrammatical subject-verb number trials are reduced when an intervening attractor agrees in number with the verb (Tanner et al., 2014, 2017). Second, Ryu and Lewis (2021) propose and demonstrate self-attention diffuseness as a proxy for degree of similarity-based interference, i.e., show that when a distractor matches the verb in number, attention is paid not only to the target, but to the distractor as well (see also Ryu & Lewis, 2025; Timkey & Linzen, 2023). The joint consideration of these results then leads to the association of attention diffuseness with the P600, at least in the specific case of ungrammatical number agreement attraction sentences.

While the fact is immediately apparent that the naturalistic storybook stimulus used here consists of grammatical material rather than ungrammatical material, it is the case, however, that cue-based retrieval theory also makes the prediction that sentences like (3b) are more difficult than sentences like (3a; examples adapted from Bock and Miller, 1991, p. 45).

(3) a. The cost 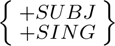 of the improvements 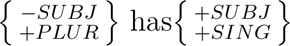 not yet been estimated.

b. The cost 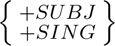 of the improvement 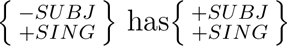 not yet been estimated.

In both (3a) as well as (3b), the {+*SUBJ*} and {+*SING*} cues for the verb *has* match perfectly the features for the subject *cost*. In (3b), but not (3a), however, *improvement* also has the {+*SING*} feature, making it more difficult for *has* to retrieve the correct target. This is known as inhibitory interference, and like facilitatory interference, we will not here get into the possible cognitive-psychological causal details, although the reader is directed to Vasishth and Engelmann (2021).

As compared to facilitatory interference in ungrammatical sentences, experimentally, inhibitory interference in grammatical trials is less borne out (see Jäger et al., 2017; Vasishth & Engelmann, 2021), with some studies failing to observe inhibitory interference (Dillon et al., 2013; Lago et al., 2015; Tanner et al., 2014, 2017; Wagers et al., 2009) — and thus labeling agreement attraction an asymmetrical phenomenon — but others indeed observing inhibitory interference effects (Franck et al., 2015; Lee & Garnsey, 2015; Martin et al., 2012, 2014). Concerned, however, that previous interference studies were under powered, Nicenboim et al. (2018) conduct a high-powered investigation and indeed do find evidence for inhibitory interference.

Important for the present project is the fact that the proposal of Ryu and Lewis (2021) — i.e., of attention diffuseness as a proxy for degree of similarity-based interference — extends to inhibitory interference, as well. Thus, we now take one step forward — from ungrammatical stimuli to grammatical stimuli — to associate (normalized) attention entropy with electrophysiological effects. Martin et al. (2012) analyze EEG responses to Spanish sentences in which a noun must be retrieved as antecedent of a nominal-ellipsis-allowing determiner, while an inaccessible noun intervenes:

(4) a. Marta se compró la camiseta{+*F EM* } que estaba al lado de la falda{+*F EM* } y Miren cogió otra{+*F EM* } _ para salir de fiesta.

‘Marta bought the t-shirt{+*F EM* } that was next to the skirt{+*F EM* } and Miren took another{+*F EM* } _ to go to the party.’

b. Marta se compró la camiseta{+*F EM* } que estaba al lado de la vestido{+*M ASC*} y Miren cogió otra{+*F EM* } _ para salir de fiesta.

‘Marta bought the t-shirt{+*F EM* } that was next to the dress{+*MASC*} and Miren took another{+*F EM* } _ to go to the party.’

Analyzing all electrodes, and in a 400–1000 ms time window after critical word onset, they find more positive *µ*V when the attractor is the same grammatical gender as the determiner (4a), then when it is not (4b). Martin et al. (2014) perform a follow-up study using similarly structured, but object-extracted relative clause stimuli. They find: 1) greater negativity in anterior channels 100–400 ms after critical word onset when the attractor is the same gender as the determiner (cf. Terpstra et al., 2025 and Fig. 7); and 2) a larger P600 in posterior channels 1200–1400 ms after critical (700–900 ms after post-critical) word onset when the attractor was different in gender than the determiner. Two of these three electrophysiological results pattern with the operating hypothesis that the presence of a gender-agreeing distractor leads to more diffuse attention (greater NAE) and greater difficulty retrieving the target antecedent, and that is observed as greater deviance in the electrophysiological signal compared to the non-interference condition. Indeed, this is what is indicated in the clusters in Fig. 7: increases in GBA NAE associated with beta coefficients (both positive and negative) of greater magnitude linking to signed source data.

While the EEG results of Tanner et al. (2014, 2017) are only indirectly applicable to the present investigation, and while the EEG results of Martin et al. (2012, 2014) are murky, they both, in conjunction with the computational modeling work connecting diffuseness of self-attention to similarity based interference during cue-based memory retrieval (Ryu & Lewis, 2021, 2025; Timkey & Linzen, 2023), support a preliminary association between normalized attention entropy and the electrophysiological results observed here. An important point that has not yet been raised, however, and which also makes direct comparison more difficult, is the fact that the discussed similarity-based interference studies involve either nominal retrieval for a subject or nominal retrieval for an antecedent — only two types linguistic dependency. This is in stark contrast to the 23 distinct Universal Dependency dependency types covered by the aggregate mixed GBA model. Thus, while the present work makes an interesting first pass, much more work will be needed in order to flesh out the details.

We end this portion of the discussion on a pair of notes. The first is that the idea of attention entropy as retrieval difficulty bears some resemblance to the notion of referential ambiguity (Nieuwland & Van Berkum, 2006, 2008; Nieuwland et al., 2007; Van Berkum et al., 1999, 2003, 2007, inter alia). Similar to how attention cannot be concentrated entirely on the true target in the presence of a distractor, in the case of referential ambiguity, a discourse entity cannot be uniquely selected from the situation model. While the (sustained) frontal negative shift (Nref; referentially induced frontal negativity) EEG component associated with this phenomenon onsets ∼300 ms, the shift can be sustained through 900 (Van Berkum et al., 2003) or even 1500 ms (Nieuwland & Van Berkum, 2006; Nieuwland et al., 2007) post-stimulus onset, thus bringing it closer in range of the late effects identified here. While this phenomenon and framework would be an interesting domain for future computational modeling, it is not a likely interpretation of the MEG effects found for normalized attention entropy for three reasons: 1) if anything, the temporal response functions shown in Fig. 7 are biphasic, and certainly not sustained shifts; 2) the GBA cue-based retrieval model from which NAE is derived considers only the current sentence; i.e., it has no knowledge of or even ability to consider the larger, preceding, situational context; and 3) the Universal Dependencies framework underlying the GBA does not consider anaphora, i.e., assign dependencies or draw arcs between coreferring entities.

Lastly, recent work by Brouwer and colleagues (Aurnhammer et al., 2023; Brouwer & Hoeks, 2013; Brouwer et al., 2012, 2017; Delogu et al., 2019, 2021, 2025, inter alia) has developed the Retrieval-Integration (RI) theory of language processing. Seeking a parsimonious account for “semantic” P600 effects like those observed for stimuli such as *The hearty meal was devouring…* (Kim & Osterhout, 2005, p. 207), the RI framework recasts N400 and P600 amplitude as indexing: context-sensitive, word meaning retrieval from long-term memory; and new information integration for mental representation update, respectively. The RI account states that every word should be associated with a biphasic N400-P600 complex for lexical activation/retrieval, and subsequent integration/update (Delogu et al., 2025). Indeed, by annotating surprisal and NAE for all applicable words — 100.0% and 37 .6%, respectively — and then observing N400- and P600-like effects, respectively, the present results are quite consistent with the RI proposal.

### Conclusion

The present project demonstrates that, even in the age of ever larger and more powerful artificial neural network models for language, there still remains a place in cognitive neuroscience for interpretable, human-bias-informed models of sentence comprehension. Our work extends recent behavioral investigations into attention diffuseness as similarity-based interference to the neural domain, and contributes to an ever-growing foundation for understanding how algorithmic-level proposals of sentence processing map onto implementation-level neural mechanisms.

The temporal dissociation observed for disambiguation and memory retrieval provides crucial evidence that these theoretically motivated cognitive processes have distinct neural substrates, supporting their treatment as separate components within algorithmic theories of sentence processing. Moreover, by demonstrating that interpretable neurocomputational models can achieve meaningful brain alignment while maintaining explanatory transparency, this work offers a methodological blueprint for bridging the gap between algorithmic proposals and neural implementation. Overall, this research contributes to the unification of disambiguation and memory retrieval theories of human sentence processing, opening up a number of interesting future directions to develop better, more interpretable, more neurologically plausible models of the systems that power the human mind.

A couple of directions present themselves as interesting avenues for future research. The first involves integrating disambiguation and memory retrieval into a single unified model. This follows clearly in the footsteps of Lewis and Vasishth (2005), who construe left-corner phrase structure parsing as cue-based memory retrieval. Benefits to starting from a neural network-parameterized transition-based dependency parser and using parse state embedding representations for retrieval calculation, however, include: 1) the maintenance of the ability to calculate disambiguation metrics over beams of parse states; and 2) broad coverage applicability for free — i.e., applicability to arbitrary (grammatical) language like that found in audiobook stories. As has already been an undercurrent in the present project, incremental dependency parsing and cue-based memory retrieval pose an elegant pairing, with retrieval triggered whenever a dependency is assigned by the parser.

The second direction — which could either be integrated into a unified model or implemented separately — involves properly accounting for degree of disambiguation work attributable to syntax — the surprisal measure calculated in the present project being lexical in nature and only syntactially informed (see Franzluebbers et al., 2024, for an example of decomposing surprisal from a generative dependency parser into lexical and syntactic components, however). Indeed, several recent proposals have attempted to get at this quantity by calculating the dissimilarity, incompatibility, or divergence between probability distributions over syntactic analyses (Wang et al., 2024, 2025; Dunagan, 2025, Ch. 3), the idea being that a large change in syntactic analysis distribution is associated with a large degree of syntactic processing work.

## Footnotes

1 We presently set aside consideration of sentence processing following the “parallel” presentation of a complete, sentence-length stimulus, although the reader is referred to e.g., Asano and Yokosawa (2011), Dunagan et al. (2025), Massol and Grainger (2025).

2 With this statement, we set aside state-of-the-art models which receive mid-training, post-training, instruction fine-tuning, reinforcement learning from human feedback, reinforcement learning from verifiable rewards, etc., and pick out this specific class of large language model.

3 This incremental nature is in contrast to both bidirectional masked language modeling, as well as the parsing scenario in which the entirety of the input string is made available from the onset, e.g., graph-based parsing.

4 The cluster-forming *p*-threshold is a parameter that controls cluster size in space, rather than determining statistical significance.

5 While 10,000 permutations is the accepted standard for cluster-based permutation testing in electrophysiological data analysis, the number of permutations is limited by the number of participants — with 12 participants, the procedure is limited to 4,095 possible permutations.

6 We set aside agreement attraction in production (“proximity concord,” Quirk et al., 1972; “attraction error,” Zandvoort, 1961), and limit attention strictly to comprehension, although the reader is directed toward the extensive literature in the domain of production (Bock & Cutting, 1992; Bock & Eberhard, 1993; Bock & Miller, 1991; Eberhard, 1997; Eberhard et al., 2005; Fayol et al., 1994; Hartsuiker et al., 2003, inter alia).

